# LincRNAs enable germ cells differentiation by promoting PUF proteins condensation

**DOI:** 10.1101/2023.08.27.554978

**Authors:** Roni Falk, Noa Gilad, Hanna Achache, Yisrael Rappaport, Reut Shabtai, Hasan Ishtayeh, Laura Wolovelsky, Yonatan B. Tzur

## Abstract

Successful tissue homeostasis depends on a balance between proliferation and differentiation. Two PUF proteins, FBF-1 and FBF-2, maintain stem-cells proliferation in *C. elegans* germ-cells by binding and destabilizing transcripts which promote meiotic entry. However, it is unclear how meiosis initiates because the FBF are also present at significant levels in late proliferative and early meiotic cells. We found that the three long-intergenic-non-coding RNAs (lincRNAs) that bind the FBF proteins promote timely meiotic entry. Deletion of the lincRNA genes leads to additive reduction in progenitor cell number and fertility. In the lincRNAs deletion mutant, expression of many known FBF-2 targets is significantly lower, suggesting over-activation of FBF-2. In this mutant, FBF-2 localization in perinuclear condensates is reduced, and its cytoplasmic fraction increases. Moreover, FBF-2 association with the germline P-granules decreases without the lincRNAs. Our results indicate that lincRNAs act to promote meiotic differentiation by spatially restricting pro-proliferation factors into phase-separated granules.

## Introduction

In many developmental processes, stem cell identity is maintained in a specialized niche through intercellular signaling. As cells physically move away from the niche, they stop proliferating and initiate the differentiation program. This occurs during gametogenesis in many metazoans *e.g.*, mammalian spermatogenesis (reviewed in ^1^). Optimal fertility relies on careful balance between mitotic cell cycles, which lead to cellular proliferation, and entry into meiosis, which initiates gamete differentiation ^2^. Partial or complete infertility are caused by genetic mutations that lead to either a premature or delayed onset of differentiation (*e.g., fbf-1 fbf-*2 and *gld-2 gld-*1 in *C. elegans, Pumilio* and *Bam* in *D. melanogaster*, and *Pum1* and *Cyp26b1* in mice) ^3–10^. One of the best studied examples of this regulated transition from mitosis to meiosis occurs in *C. elegans* gonads, where germ cells are spatio-temporally organized from stem cells distally to mature oocytes proximally (Fig. 1a, top). Here we focus on germ cells in the distal gonad as they move through the Progenitor Zone (PZ) into Leptotene/Zygotene Zone (LZ) (Fig 1a, bottom). The more distal “early PZ” includes germline stem cells, while the more proximal “late PZ” includes cells that undergoes mitotic cell cycles as well as meiotic S-phase cells ^11^. As germ cells move from the PZ to the LZ, they enter the first stages of meiotic prophase I. The niche for germline stem cells is formed by a single somatic cell, the distal tip cell, which signals via the GLP-1/Notch pathway to germ cells in the early PZ (Fig 1a, reviewed in ^11–13^).

**Figure 1:**
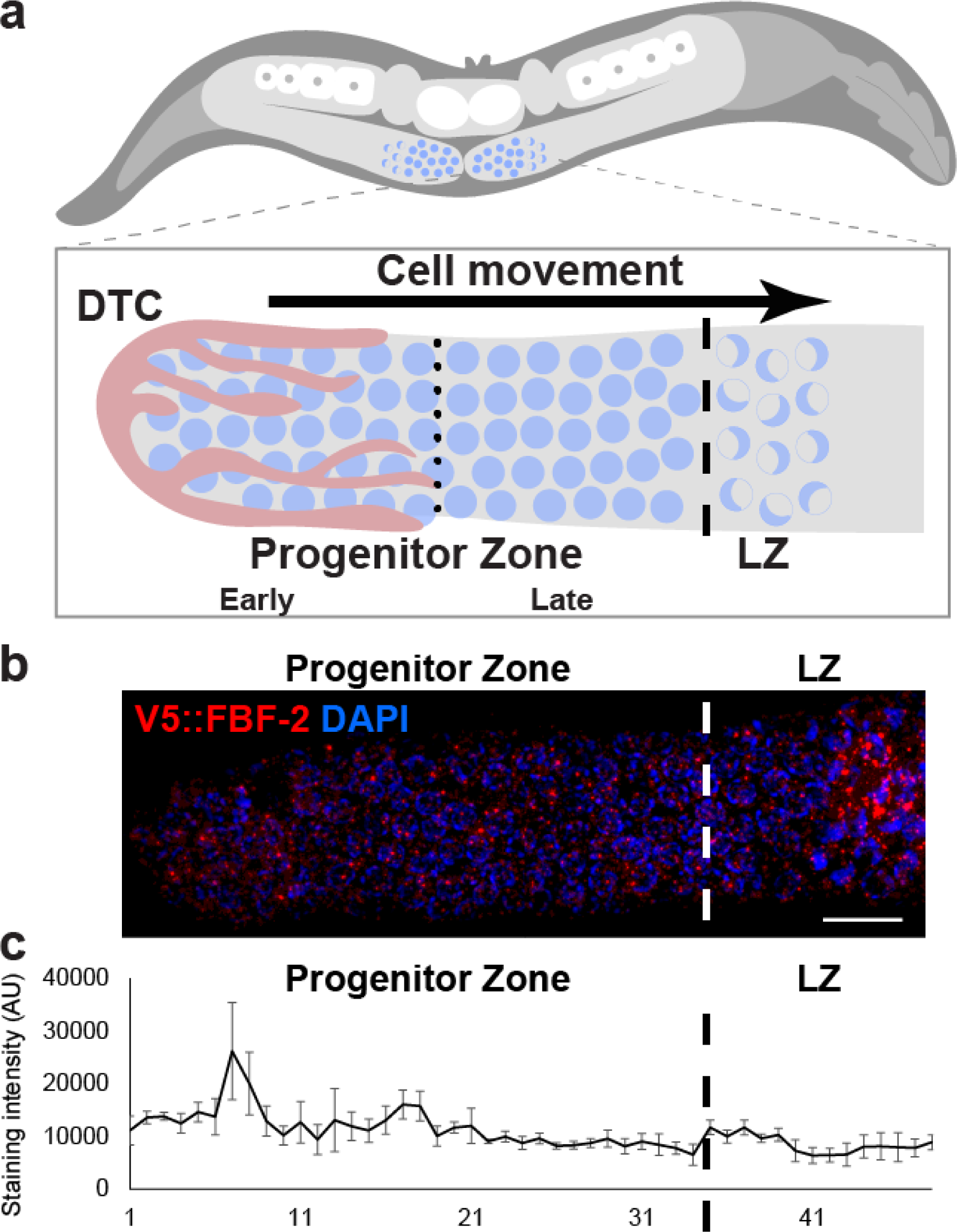
FBF-2 levels are similar in progenitor zone and early differentiation stages in the *C. elegans* gonad. a. Illustration of the *C. elegans* gonad distal side. The distal tip cell (DTC) is marked in pink. b. Distal side of a gonad stained with V5::FBF-2 (red) and DAPI (blue). c. Quantification of the average level of V5::FBF-2 along the distal side of the gonad. Dotted line indicates the border between early and late progenitor zone. Dashed line indicates the border between progenitor zone and early meiotic stages. DTC: distal tip cell, LZ: Leptotene/Zygotene, N value = 3, Scale bar = 10 μM.

Key regulators of mitotic proliferation are FBF-1 and FBF-2 (collectively FBF), which block meiotic entry and control the mitotic cell cycle ^4, 14–16^. These proteins are founding members of the conserved PUF (Pumilio and FBF) family of RNA-binding proteins, which control development from worms to humans ^17^. FBF-1 and FBF-2 each binds thousands of mRNAs, reduce their stability and\or repress their translation ^18, 19^. They also bind a few long intergenic non-coding RNAs (see below) ^18, 20^.

FBF-1 and FBF-2 are 89% identical in their amino acid sequence and largely bind the same transcripts ^18, 20^. While single mutants are fertile, double mutants have small germlines composed only of differentiated sperm ^4^. FBF-1 and FBF-2 therefore carry-out redundant functions, but they differ in some aspects. For example, the PZ is smaller than wild-type in *fbf-1* mutants but larger than wild-type in *fbf-2* mutants ^21^. Moreover, *fbf-1* mutant germ cells move faster into meiosis, while *fbf-2* mutant germ cells move slower and have a longer cell cycle ^16^. One model that was proposed suggested that the level of the FBF drops as germ cell progress through the PZ, thus relieving their repression on meiosis promoting transcripts (reviewed in ^22^). Yet, FBF-1 and FBF-2 are both still present in meiotic S-phase cells in late PZ and early meiocytes in the LZ ^19, 21, 23^. Therefore, although FBF maintain proliferation in the early PZ, how that activity is turned down in the late PZ and LZ so that cells enter meiosis is largely unknown.

Long intergenic non-coding RNAs (lincRNAs) are classified by a length longer than 200 nucleotides and transcription from loci that do not code for proteins ^24^. They are usually produced by RNA polymerase II, capped, and polyadenylated (reviewed in ^25–34^). Long non-coding RNAs were previously shown to restrict the action of PUF proteins in mammalian cell culture ^35, 36^, and in worms three lincRNAs bind the FBF ^18, 20^. We therefore hypothesized that lincRNAs may be involved in FBF inhibition.

Here we show that the removal of all three lincRNAs known to bind the FBF display an additive loss of fertility and a reduction in the number of PZ cells. Deletion of those genes reduces the levels of FBF-2 bound transcripts, suggesting the lincRNAs play a role in repressing FBF-2. We found that FBF-2 perinuclear condensation increases as the cells exit the niche in a lincRNA dependent manner. This indicates that the lincRNAs act to restrict FBF-2 mRNA repression by promoting its condensations and thus enable proper meiotic entry.

## Results

### FBF-1 and FBF-2 are expressed in early meiotic prophase

According to the accepted model, within the progenitor zone (PZ) of *C. elegans* gonads, cell entry into meiosis is prevented by repression of key meiotic transcripts by the FBF proteins. Nevertheless, FBF proteins have also been reported in germ cells in leptotene/zygotene (LZ) ^19, 21^. Chen et al. used a strain with an N-terminal V5 tag inserted at the genomic locus of *fbf-2,* to quantify the relative levels of FBF-2 at the distal part of the gonad of adults eight hours post L4/adult molt ^23^. We repeated this analysis under our experimental conditions (20-24 hours post L4 stage, see methods) and, in agreement with Chen et al., we found that FBF-2 levels reach a peak, 7-8 nuclei rows from the distal end and then quickly drops to ∼33% of its peak level (Fig. 1b-c). Surprisingly, we found that during the first meiotic stage, Leptotene/Zygotene (LZ), the level of FBF-2 is very similar to the level measured at the PZ after the sharp early peak (Fig 1b-c). In line with previous results, we found that FBF-1 expression was mostly stable throughout the PZ but dropped to 60% at the first LZ nuclei (Fig. S1). Our analyses indicate that in late PZ nuclei the levels of both FBF-1 and FBF-2 are as high as most of the early PZ nuclei. At the entry to meiosis FBF-2 levels are maintained and FBF-1 levels drop, but it is still present in considerable amount. Taken together, we draw two possible conclusions. Either meiosis is initiated in the late PZ despite the presence of FBFs in their repressive state, or some mechanism inhibits their action post translationally in late PZ to enable meiotic entry. We hypothesized that lincRNAs are involved in promoting the latter.

### Three FBF-bound lincRNAs have additive roles in fertility

We previously reported the engineering of six *C. elegans* worm strains with genomic deletions of lincRNA genes that are highly expressed in the gonad. We did not detect a considerable reduction in fertility in any of them ^37^. Therein, we suggested that the lack of significant fertility loss could stem from redundant roles of these genes. Two of these lincRNAs, *linc-4* and *linc-7,* were previously shown to bind the FBF proteins ^18, 20^, and their deletion led to a very slight reduction in brood size (Fig. 2a and 37). To find if the two lincRNAs act redundantly, we quantified the brood size of the double mutant, and found it has significantly smaller progeny than wild-type worms as well as both the single mutants (Fig. 2a).

**Figure 2:**
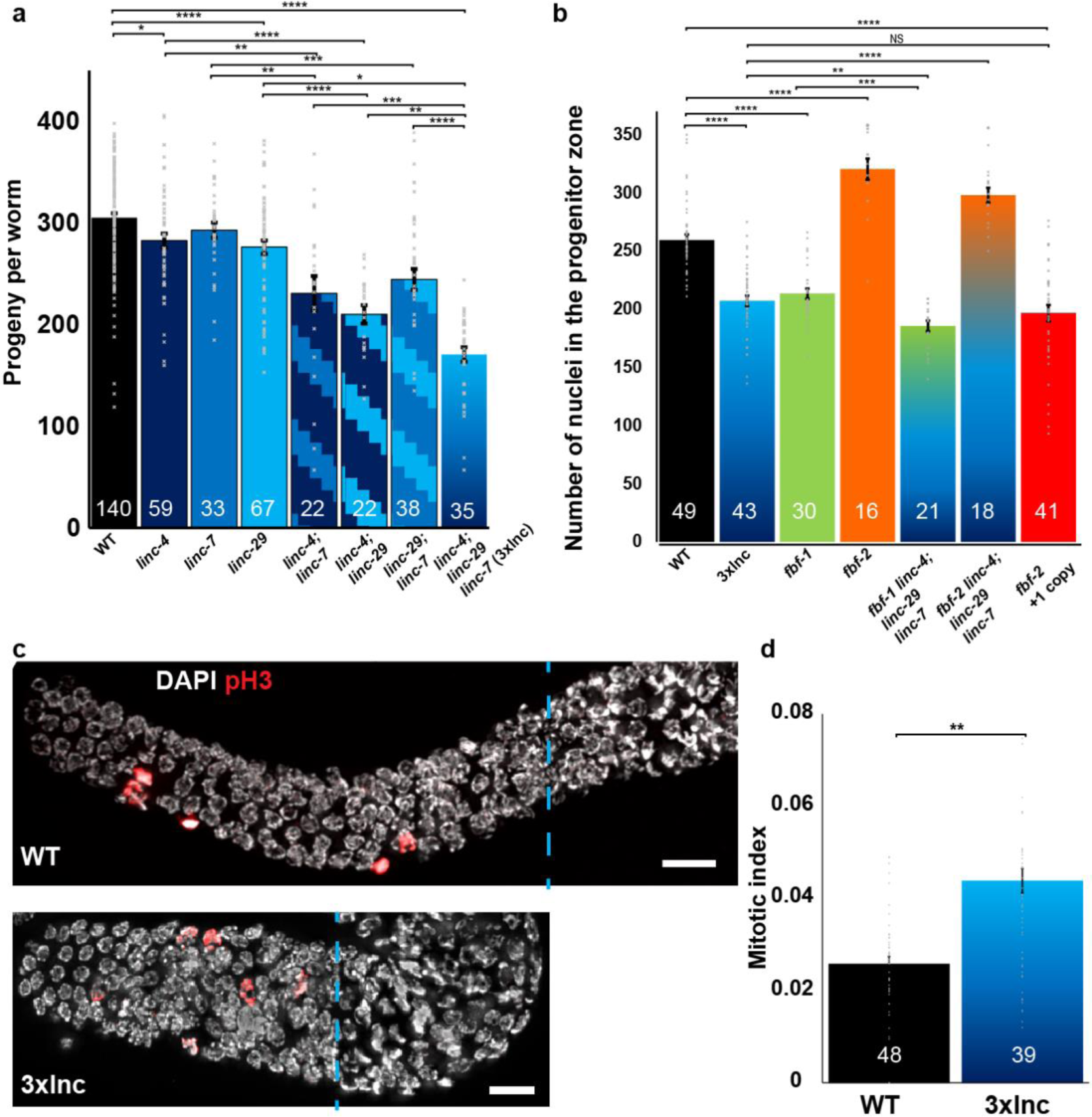
Deletion of three lincRNAs reduce fertility and the number of progenitor zone nuclei. a. Average self-progeny number of the indicated genotypes. b. Average number of nuclei in the progenitor zone in gonads of the indicated genotypes. c. Distal side of wild-type (top) and 3xlnc (bottom) gonads stained with pH3 (red) and DAPI (white). d. Average mitotic index of wild type and 3xlnc. Dashed lines indicate the border between progenitor zone and early meiotic stages. N values are marked on the columns. Mann Whitney *p* value: *<0.05, **<0.01, ***<0.001, ****<0.0001. Scale bar = 10 μM.

The additive effect on fertility of the two FBF binding lincRNAs supports the hypothesis that the lincRNAs play a part in the meiosis entry tipping point within the FBF pathway. In line with this possibility, the levels of *linc-4* and *linc-7* increase from the early PZ to the late PZ, as well as from late PZ into the LZ and early meiosis ^37, 38^ (Fig. S2). The only other lincRNA known to bind the FBF in hermaphrodites is *linc-*29 ^18, 20^. Similar to *linc-4* and *linc-7,* a complete deletion of the *linc-29* gene leads to a small reduction in brood size (Fig. 2a). The double mutants of *linc-4; linc-29* and *linc-29; linc-7* had significantly smaller progeny than both wild-type and single mutant worms. In the triple *linc-4; linc-29; linc-7* strain (hence 3xlnc) the brood was 30% lower than wild-type worms and significantly lower than the double mutants (Fig. 2a). We verified that this effect stems from *linc-29* transcripts and not from off-target effects, or unrelated deletion of a DNA element, by using RNA silencing to reduce the expression of *linc-29.* RNAi of *linc-29* significantly reduced the progeny size compared to control RNAi (Fig. S3a), indicating that the fertility effect is dependent on *linc-29* transcripts.

To test if the effect of lincRNAs was dependent on the binding of the lincRNAs to the FBF we aimed to create a minimal mutation that will disrupt this binding. Among the three lincRNAs, *linc-7* has the longest iClip peak withing the FBFs, yet it contains only 1 characteristic UGUNNNAU FBF binding element (FBE) ^18, 20^. We therefore engineered a strain with a genomic deletion of the *linc-7* FBE (*linc-7ΔFBE). Linc-4; linc-29*; *linc-7ΔFBE* triple mutant worms displayed a significant reduction in brood size compared to *linc-4; linc-29* worms. The progeny size of *linc-4; linc-29; linc-7ΔFBE* was very similar to the 3xlnc (*linc-4; linc-29; linc-7*) strain (Fig. S3b). This indicates that correct binding of the FBF to *linc-7* is required for optimal fertility. Taken together these results show that the three lincRNAs have additive roles in fertility by binding to the FBF.

### The lincRNAs play roles in normal entry to meiosis

FBF-1 and FBF-2 negatively regulate each other and were suggested to have opposite effects on mitotic cell cycles ^16, 21^. Consistent with that idea, the *fbf-1* mutant PZ has fewer nuclei than wild-type, while the *fbf-2* mutant PZ has more nuclei (^16^ and Fig. 2b). To find if the 3xlnc mutant affects the number of PZ nuclei or mitotic index, we stained wholemount gonads with DAPI and pH3 antibodies to mark M-phase nuclei (Fig. 2c). The 3xlnc mutant PZ was significantly smaller than wild-type with fewer nuclei (Fig 2b), but its mitotic index was significantly higher than wild-type (Fig. 2d). The 3xlnc effect is thus similar to *fbf-1* and opposite to *fbf-2* (Fig. 2b ^16^). To further test the genetic interactions between the lincRNAs and the *fbf*, we analyzed the effect on the PZ when the lincRNA genes are removed together with mutants in *fbf* genes. We found that *fbf-1 linc-4; linc-29; linc-7* worms have significantly less nuclei in the PZ than 3xlnc and *fbf-1* mutants (Fig. 2b). This suggests that the effects of removal of the lincRNAs and *fbf-1* work in parallel. In contrast, the proliferative population of *fbf-2 linc-4; linc-29; linc-7* is larger than wild-type worms and almost as high as that of *fbf-2* worms (Fig. 2b). Therefore, the reduction in the PZ population observed when the lincRNAs are removed depends, at least in part, on the presence of *fbf-2*.

Importantly, the similarities between the reduced PZ population observed in *fbf-1* and 3xlnc, in contrast to *fbf-*2, support a model by which the lincRNAs act to repress FBF-2. To test this possibility, we quantified the PZ nuclei number in a strain with an extra copy of *fbf-2* (YBT103, see Methods). In this strain we found smaller PZ than wild-type worms which was practically identical to 3xlnc (Fig. 2b). Not surprisingly, the smaller PZ population we found in the presence of ectopic expression of *fbf-2* is opposite to the larger PZ observed in *fbf-2* mutant. Taken together these results support the hypothesis that the lincRNAs act to inhibit the roles of FBF-2. We therefore focused our efforts on analyzing the interactions between the lincRNAs and FBF-2.

### Deletion of three lincRNAs leads to reduced levels of FBF-2 bound transcripts

FBF-2 translationally represses its bound transcripts ^19^. To test whether FBF-2 also reduces their stability, we used RNA-Seq analysis to compare mRNA levels in wild-type vs *fbf-2*. We found that 1303 genes were significantly upregulated in *fbf-2* (Fig. S4, Table S1a). Among these, 254 were previously found to bind FBF-2 ^20^ (*p* value = 2.1*10^−15^, by the by Fisher Exact test). We also found 1359 genes downregulated in *fbf-2*, but among these, only 34 were FBF-2 bound mRNAs (Fig. S4, *p* value ∼ 1, by the Fisher Exact test). Together, these analyses suggest that FBF-2 regulates stability of some mRNAs.

The reduced PZ population observed in 3xlnc is opposite to the increased population observed in *fbf-2,* and similar to worms with ectopic expression of *fbf-2*. This suggests a model by which the lincRNAs act by binding and repressing FBF-2. The model predicts that in 3xlnc, where the lincRNAs are missing, FBF-2 will be overactive and the levels of the transcripts it binds and represses will decrease.

To test our prediction, we analyzed the changes in the transcriptome of 3xlnc compared to wild type by RNA-Seq. We found that in 3xlnc 494 genes were significantly downregulated compared to wild type. 141 genes within this group were also known FBF-2 bound genes (Fig. 3a, S4, p value < 2.3*10^−24^, by the Fisher Exact test). As a control, we analyzed genes that were upregulated in 3xlnc. We found that 1290 genes were significantly upregulated in 3xlnc vs WT. Only 14 of the upregulated genes were known FBF-2 bound genes (*p* value ∼1, by the Fisher Exact test).

**Figure 3:**
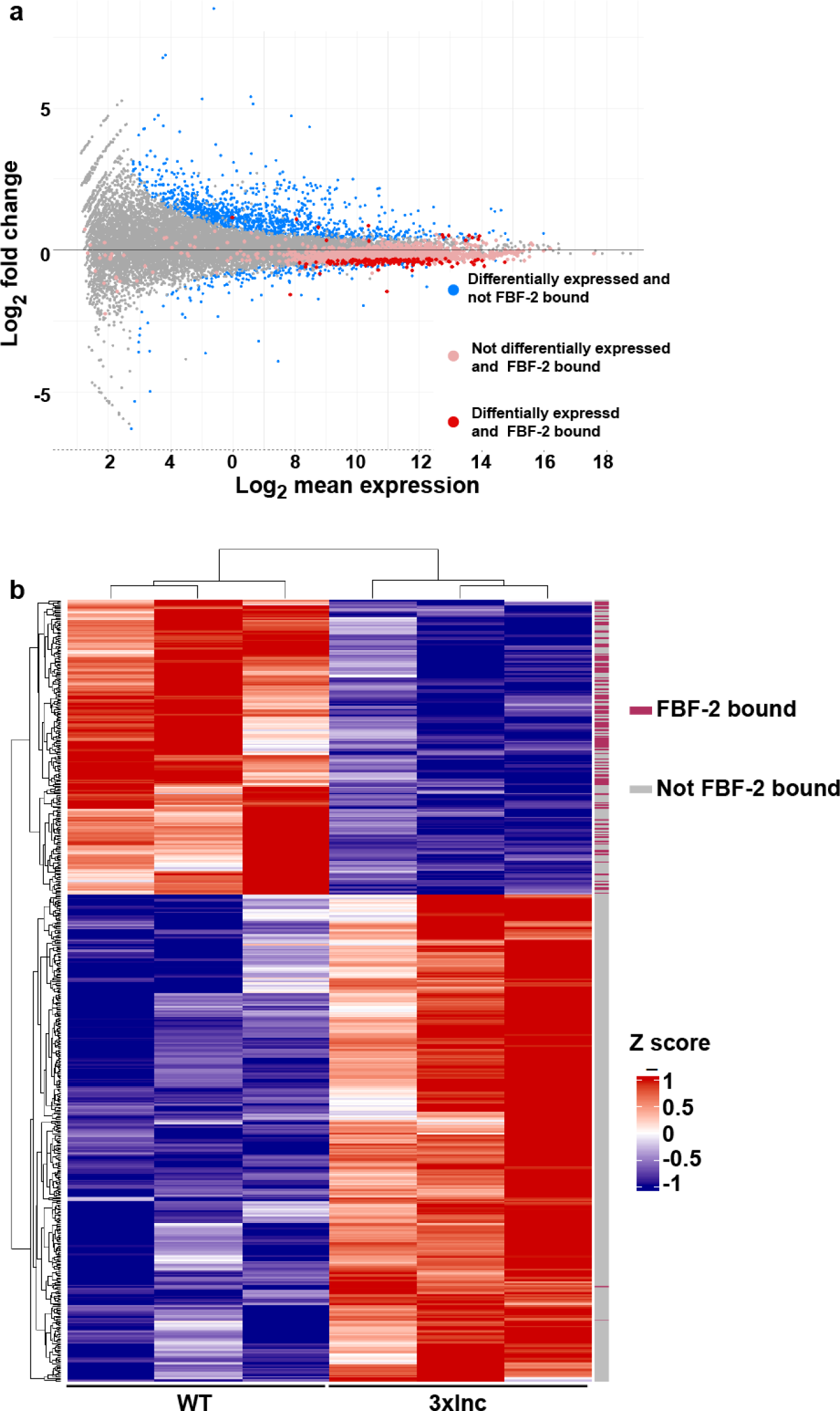
FBF-2 bound genes are downregulated without the lincRNAs. a. Expression differences in 3xlnc vs. wild type for each gene plotted against the Log_2_ average expression. The x axis is the DESeq2-calculated baseMean, and the y axis is the DESeq2-calculated lfcMLE (log2 of the fold-change). Differentially expressed genes are marked in blue, FBF-2 bound genes which are not significantly differentially expressed are marked in pink and FBF-2 bound genes which are significantly differentially expressed are marked in red. Note that most FBF-2 bound genes are downregulated in 3xlnc. b. Clustered heat map representation of germline enriched genes ^39^ which are differentially expressed, with FBF-2 bound boolean annotations (crimson) ^20^. Genes are clustered according to their expression profile. Normalized expression values were standardized according to their z-score and then clustered by hierarchical clustering.

**Figure 4:**
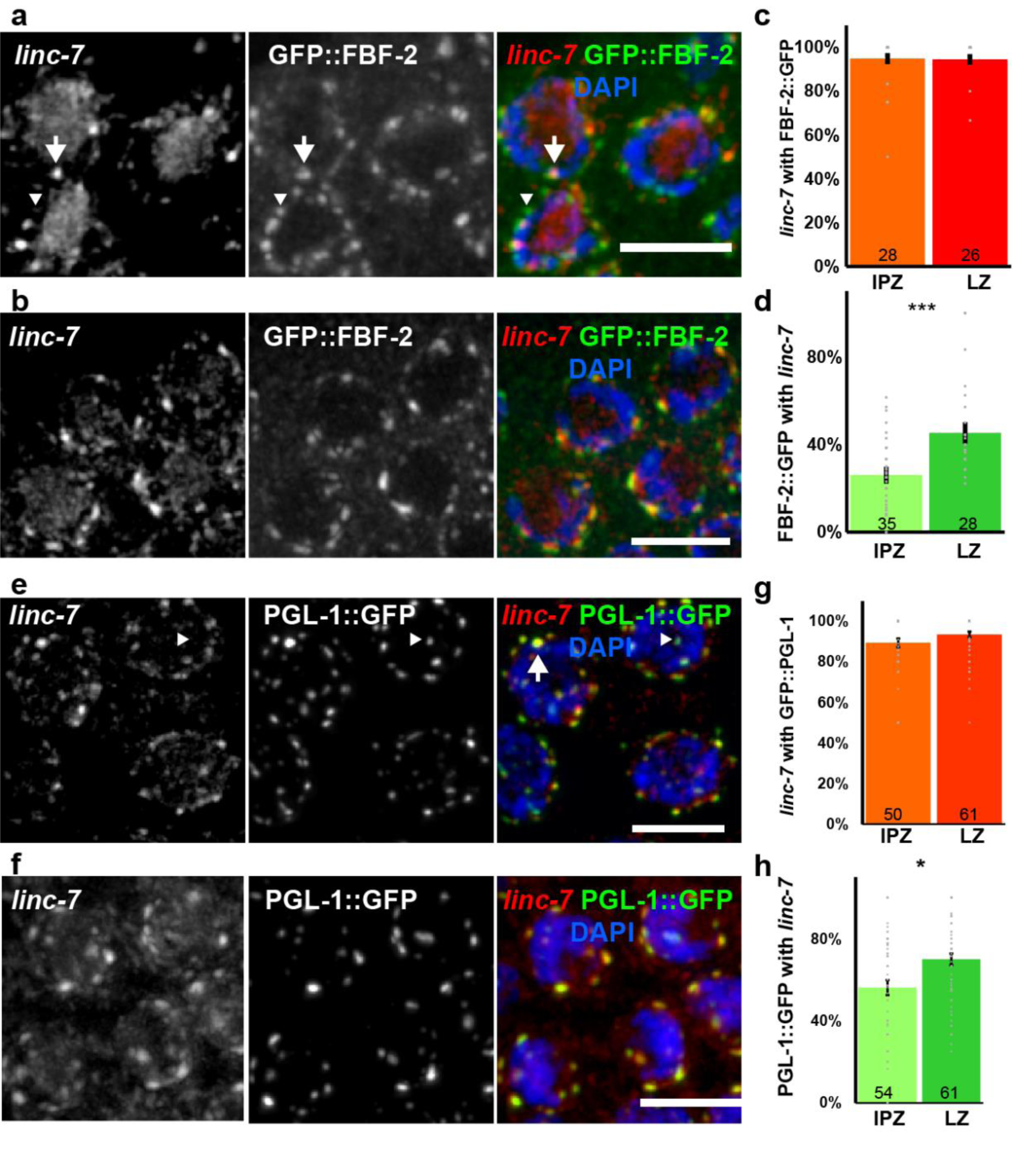
*Linc-7* is localized to perinuclear foci with FBF-2 and PGL-1. a-b. Late progenitor zone (a) and LZ (b) nuclei stained with *linc-7* smFISH (red), GFP::FBF-2 (green) and DAPI (blue). c. Average percentage of *linc-7* perinuclear foci that co-localize with GFP::FBF-2. d. Average percentage of GFP::FBF-2 perinuclear granules that co-localize with *linc-7*. e-f. Late progenitor zone (e) and LZ (f) nuclei stained with *linc-7* smFISH (red), PGL-1::GFP (green) and DAPI (blue). g. Average percentage of *linc-7* perinuclear foci that colocalize with PGL-1::GFP. h. Average percentage of PGL-1::GFP perinuclear condensates that colocalize with *linc-7*. N values are marked on the columns. Scale bar = 5 μM. Arrow: colocalized foci. Arrowhead: protein focus that do not co-localize with *linc-7.* Mann Whitney *p* value: *<0.05, ****<0.0001. lPZ: late progenitor zone. LZ: Leptotene/Zygotene.

To focus on the effect of the removal of the lincRNAs on fertility, we analyzed only germline enriched genes ^39^. We found that among the 2900 germline genes, 367 were significantly upregulated in 3xlnc but only two were known FBF-2 bound genes. Remarkably, of the 221 germline enriched genes which were significantly downregulated in 3xlnc, 102 were known FBF-2 bound genes (Fig. 3b, p value < 4.4*10^−12^, by the Fisher exact test). Our results indicate opposite effects on FBF-2 bound transcripts between *fbf-2* vs the removal of the lincRNAs. This observation supports the prediction that FBF mediated mRNA repression is overactivated without the three lincRNAs, in line with our model. Taken together, our analyses indicate that the combined expression of the three lincRNAs act to inhibit the FBF-2 mediated destabilization of germline transcripts.

### *Linc-7* partially co-localize with FBF-2 and P granules

To gain insights into the mechanism of action of the lincRNAs, we sought to determine their cellular localization. For this analysis, we used single molecule fluorescent in situ hybridization (smFISH) and focused on *linc-7.* The other two *linc* RNAs, *linc-4* and *linc-29,* are shorter (216 and 629 nucleotides respectively compared to 1130 nucleotides for *linc-7*), and their expression is lower (Fig. S2). The *linc-7* RNA was thus expected to produce a more robust smFISH signal. Consistent with our spatial transcriptome analysis ^37^ (Fig. S2), the *linc-7* smFISH signal increases as germ cells move from the PZ to early meiotic stages (Fig. S5a). Closer examination revealed *linc-7* in granules around the outer nuclear envelope, as well as at lower levels within the nucleoplasm (Fig. 4). This pattern was not observed in worms with a deletion of *linc-7 (*Fig. S5b). Interestingly, *linc-7* perinuclear granule vs nucleoplasm localization becomes more pronounced in late PZ and early meiotic stages (Fig. 4a-b).

FBF-2 localizes to both the cytoplasm and perinuclear granules. (^19^ and Fig. 4a-b). We tested the possibility that FBF-2 and *linc-7* are colocalized within those perinuclear granules by staining the gonads of worms expressing GFP tagged FBF-2 with *linc-7* smFISH probes. We found that in late PZ and LZ, almost all *linc-7* granules colocalized with GFP::FBF-2 granules (95% and 98% respectively, Fig. 4a-c). Surprisingly, much lower levels of GFP::FBF-2 perinuclear granules colocalized with *linc-7* granules, yet the level of colocalization significantly increased from late PZ to early meiotic stages (26±3% and 45±3% respectively, *p* value < 0.001 by the Fisher-Exact test, Fig. 4d). These results indicate that FBF-2 and *linc-7* colocalize in specific granules in cells that enter meiosis and that FBF-2 localization with *linc-7* significantly increases with meiotic entry.

FBF-2 granules were previously shown to be in partial colocalization with P granules^19^. These evolutionarily conserved ribonucleoprotein assemblies are present in the germline, mostly around the cytoplasmic side of nuclear pore complexes ^40–43^. The P granules help to maintain germline identity, as well as control mRNA stability, translation, and small RNA production (reviewed in ^44–47^). To test if the *linc-7* granules, which almost exclusively contain FBF-2, are also localized with P granules, we used a strain which expresses GFP tagged to the canonical P granule protein PGL-1 ^48, 49^. We quantified the level of localization of *linc-7* granules with P granules and found that almost all *linc-7* granules are colocalized with PGL-1::GFP granules (89% and 93% for late proliferative and L/Z respectively, Fig. 4e-g). Similar to FBF-2, the perinuclear PGL-1 granules’ colocalization with *linc-7* also increased as the nuclei proceeded from late PZ into early meiosis (56±4% and 70±3% respectively, *p* value < 0.05 by the Fisher-Exact test, Fig. 4h). Thus, *linc-7* is mostly localized to the P granules and FBF-2 perinuclear granules. Taken together, our *linc-7* localization analyses suggest that a sub-category of P granules contain both *linc-7* and FBF-2 and that the ratio of these P granules increases with meiotic entry.

### FBF-2 perinuclear condensation is reduced without the three lincRNAs

Previous reports indicated that long non-coding RNAs are involved in protein condensation ^34^. This raises the possibility that *linc-4, linc-29* and *linc-7* may play a role in FBF-2 condensation. To test whether removal of the three lincRNAs leads to a change in the localization of FBF-2, we quantified the number of FBF-2 foci in late PZ nuclei. We found a significant decrease in the average number of perinuclear granules in worms with deletions in the three lincRNAs compared to wild-type background (Fig. 5a-b, 10.7±0.7 vs 15.2±0.9 respectively, *p* value < 0.01 by the Mann-Whitney test). We found similar reduction in LZ nuclei (Fig. 5a-b, 8.4±0.5 vs 15.7±1.1 respectively, *p* value < 0.01 by the Mann-Whitney test).

**Figure 5:**
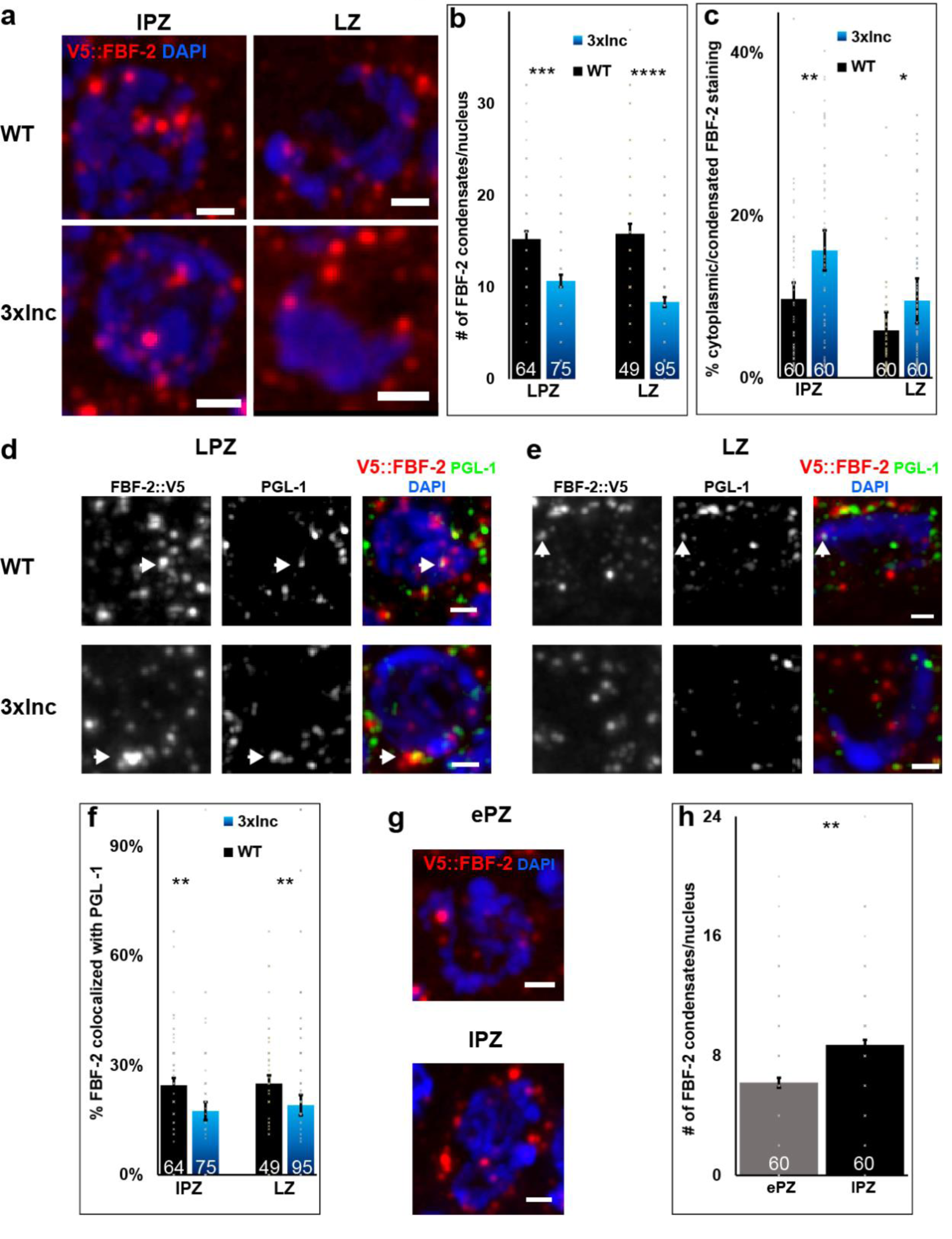
LincRNA removal reduces FBF-2 perinuclear condensation. a. Wild-type and 3xlnc late proliferative (left) and LZ (right) nuclei stained with V5::FBF-2 (red) and DAPI (blue). b. Average number per nucleus of FBF-2 perinuclear granules in wild type and 3xlnc during late progenitor zone and LZ stages. c. Average percentage of the ratio between FBF-2 staining in the cytoplasm vs nearby foci in late progenitor zone and LZ stages. d-e late progenitor zone (d) and LZ (e) nuclei stained with V5::FBF-2 (red), PGL-1 (green) and DAPI (blue). Arrow: colocalized foci. f. Average percentage of FBF-2 perinuclear foci which colocalize with PGL-1 during late progenitor zone and LZ stages. g. Early (up) and late (down) progenitor zone nuclei stained with V5::FBF-2 (red) and DAPI (blue). h. Average number per nucleus of FBF-2 perinuclear foci during early and late progenitor zone. N values are marked on the columns. ePZ: early progenitor zone. lPZ: late progenitor zone. LZ: Leptotene/Zygotene. Scale bar = 1 μM. Mann Whitney *p* value: *<0.05, **<0.01, ***<0.001, ****<0.0001.

The action of FBF-2 in inhibiting the expression of meiosis promoting genes was previously suggested to depend on the balance between the cytoplasmic and the perinuclear granules^19^. To examine the lincRNA’s role in this balance, we quantified the ratio between the level of FBF-2 staining in the cytoplasm vs a nearby perinuclear granule in each nucleus individually. We found that compared to wild type, this ratio was significantly skewed towards the cytoplasmic fraction in 3xlnc (Fig. 5c). These results suggest that the lincRNAs play a role in the condensation of FBF-2.

*Linc-7* is mainly localized in perinuclear granules which are both PGL-1 and FBF-2 positive. To test the effect of removing the three lincRNAs on the localization of FBF-2 granules within the P granules, we quantified the relative fraction of FBF-2 granules which are colocalized with PGL-1. In both late PZ and early meiotic nuclei 25% of FBF-2 perinuclear granules are colocalized with PGL-1. Conversely, in a 3xlnc background, only 19% of the FBF-2 granules were colocalized with PGL-1 (*p* value < 0.05 and 0.01 for late PZ and LZ respectively, Fig. 5d-f).

Our results point towards a connection between the perinuclear condensation of FBF-2 and the presence of the lincRNAs. To test if this connection also exists during differentiation of wild-type germ cells, we compared the number of perinuclear granules in early PZ nuclei, where the level of the lincRNAs is low, to late PZ nuclei, where the lincRNA presence increases (Fig. 5g-h, S2). We found a significant increase in the number of perinuclear FBF-2 granules as cells proceeded through the PZ (Fig. 5e-f). Taken together, these analyses indicate that the three lincRNA have roles in both condensation of FBF-2 around the nuclei and increased targeting to the P granules.

## Discussion

### LincRNAs act post-transcriptionally to repress PUF proteins

To initiate gametogenesis, germ cells in many metazoans turn off factors that promote stem cell self-renewal and delay differentiation ^13, 50, 51^. The tipping point between proliferation and differentiation must be carefully regulated to prevent aberrant outcomes that can lead to infertility. We found three lincRNAs that act during *C. elegans* oogenesis to bind and restrict the action of FBF-2 that maintains proliferation. This conclusion is supported by the rise in the lincRNA’s level as the cells progress towards meiosis (Fig. S2), the lincRNAs’ direct binding to FBF-2 ^18, 20^, the colocalization of *linc-7* with FBF-2 (Fig 4a-b), and the significant decrease in the levels of transcripts which FBF-2 restricts without the lincRNAs (Fig. 3). Not surprisingly, the increase in FBF-2 activity results in fewer progeny and fewer PZ nuclei (Fig 2). The action of the lincRNA is directly linked to FBF-2, as evident from the effect of the deletion of the FBF binding element (Fig. S3) and genetic analysis (Fig. 2).

Could the lincRNAs control FBF-2 via transcriptional control? Indeed, many lincRNAs act by altering the transcription of specific genes, most notably genes located in cis to the lncRNA gene (reviewed in ^52^). Nevertheless, we do not favor this possibility. First, among the three lincRNAs, *linc-4*’s genomic location is the closest to the *fbf* genes (∼177 kb from *fbf-*2). However, *linc-29* and *linc-7* are transcribed from different chromosomes. Second, we found no significant change in the levels of *fbf-1* or *fbf-2* RNA in 3xlnc compared to wild type (Table S1). Third, most of *linc-7* transcripts are located outside the nucleus (Fig. S5). Thus, the mechanism by which the lincRNAs reduce the FBF action is probably not due to direct control on the *fbf* transcription.

Is it possible the lincRNAs also work via FBF-1? In both 3xlnc and *fbf-1* we found similar reduction in the PZ population yet, the quadruple mutant worms show partial additive effects. (Fig. 2b). Moreover, while the progeny of the 3xlnc is dramatically lower than wild type, *fbf-1* mutation was not reported to reduce brood size (Fig. 2a). Given the minor contribution of *fbf-1* to the preservation of the PZ population, the opposite roles of the two FBF proteins, their negative effects on each other, and the similar transcript cohort they bind ^18–21, 53^, it is currently unclear which interactions exist between the lincRNAs and FBF-1.

Voronina et al. suggested that within the P granules, which interact with the nuclear pore complexes, FBF-2 associates with meiotic mRNAs and moves to the cytoplasm with the bound transcript ^19^. The results presented in this work, especially the change in the FBF-2 aggregation following the deletion of the three lincRNAs, expands this model (Fig. 6): In early PZ stages the expression of the lincRNA is low and FBF-2 can move freely between the perinuclear granules and the cytoplasm, where it destabilizes the bound mRNAs. As the nuclei move proximally, the expression of the lincRNAs rises and they bind and retain FBF-2 within P granules. This process changes the dynamics of FBF-2 localization between perinuclear granules and the cytoplasm. The retained FBF-2 has reduced ability to prevent the expression of the bound transcripts that act to initiate meiotic program, and they are eventually freed to be translated and promote differentiation. This model is supported by the specific colocalization of *linc-7* to granules that are both P granules and FBF-2 positive, the change in the balance between cytoplasmic vs perinuclear FBF-2 in 3xlnc, and the decrease in FBF-2 silencing activity at the same time the lincRNAs become more abundant.

**Figure 6:**
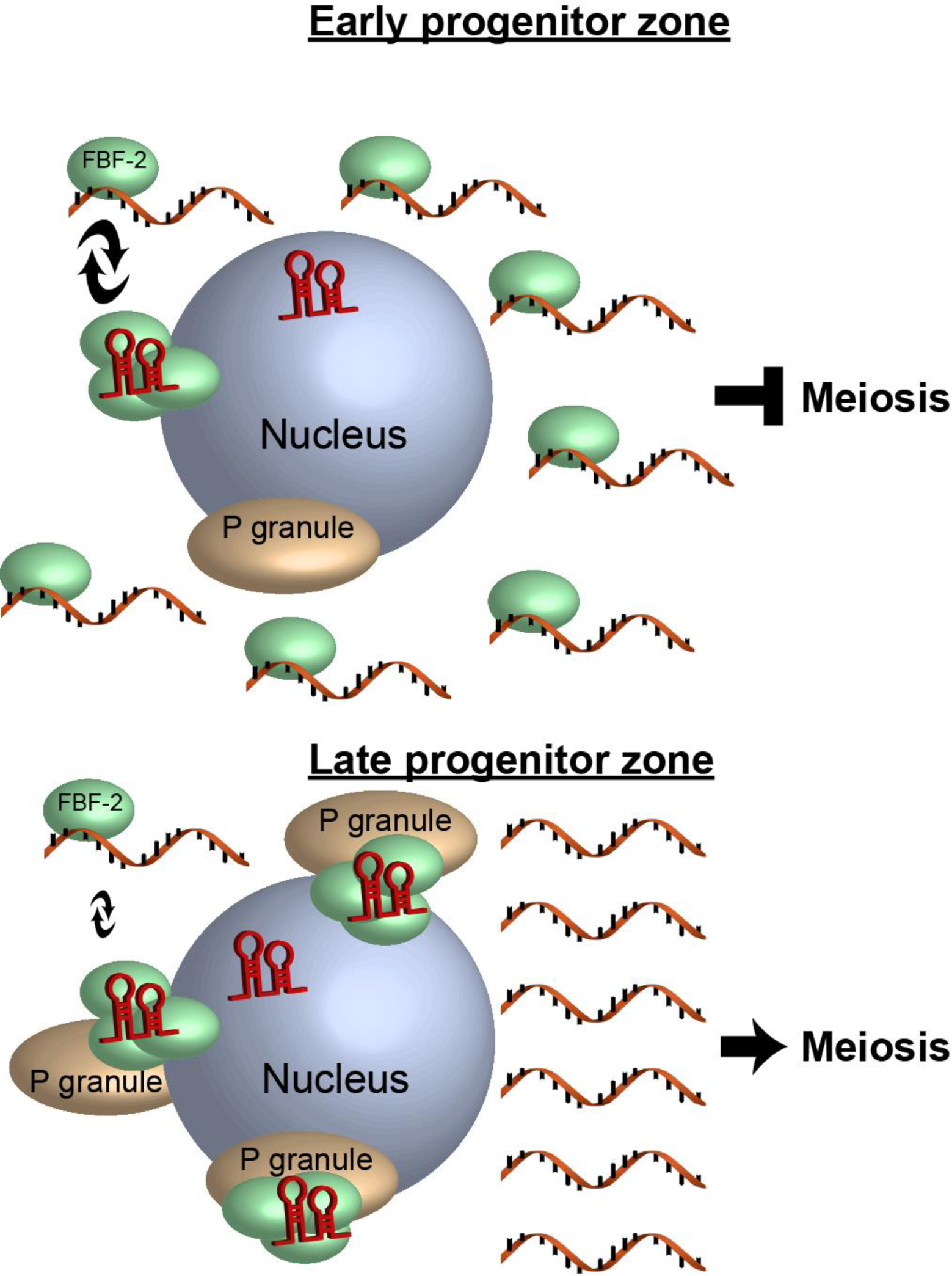
Model for the roles of the lincRNAs during meiotic entry. Illustration of the different components during proliferation of germ cells. During early stages (top) there are low levels of lincRNAs (dark red), which can bind FBF-2 (green) and promote its sequestering it to perinuclear granules. Many of the FBF-2 proteins can still shuttle back to cytoplasm where they can inhibit the expression of the meiosis promoting mRNAs (orange). During late progenitor zone (bottom), lincRNAs expression increases, they sequester more FBF-2 to perinuclear granules, together with the P granules (brown), leading to reduced FBF-2 presence in the cytoplasm. This change frees enough meiosis promoting mRNAs to initiate differentiation.

### Possible place of the three lincRNA within the framework of mitosis/meiosis decision

The triple deletion of *linc-4, linc-7,* and *linc-29* results in germline phenotypes which bear similarity to changes in *fbf* gene expression (*e.g.,* Fig. 2). However, the effect on fertility is weaker than *fbf-1 fbf-2* double mutants, in which germline proliferation halts very early, or mutants in meiosis promoting genes (e.g., *gld-1 gld-2* ^10^) where tumors form. We can envision several possibilities for the milder effect of the lincRNAs deletions. First, it is possible that the lincRNAs only affect the activity of FBF-2, resulting in partial reduction in fertility. Second, the lincRNAs may serve as a more subtle and minor arm in the meiosis entry decision, one that may become more prominent under different environmental or physiological conditions. Third, the two FBF proteins act within a negative feedback loop on each other ^19, 21, 53^, and two other PUF proteins also act to control stem cells self-renewal ^54^. Thus, within this complex PUF network, the lincRNAs may orchestrate different phases of FBF activity along several developmental stages. Fourth, as we previously predicted, lincRNAs that seem to have little or no germline roles by themselves, act redundantly with other lincRNAs ^37, 55, 56^. Indeed, we found additive roles for three of those lincRNAs. This suggests that, just as multiple PUF proteins are required to maintain proliferation, there may be more inhibitory non-coding transcripts with similar roles as the three lincRNA described here. Although *linc-4, linc-7* and *linc-29* are the only lincRNAs found to be associated with the FBF ^18, 20^, since these works were published, more lincRNAs genes were found to be transcribed in *C. elegans* ^57^. Moreover, unlike lincRNAs, data on long non-coding RNAs (lncRNAs), which are transcribed from coding regions, are lacking in *C. elegans*. The use of iClip to identify which of these lncRNAs associate with the FBFs will be challenging given these lncRNAs share sequences with coding genes. Taken together, it is very likely that along with the three lincRNAs we report here, other long non-coding RNAs may act in a similar and additive fashion to restrict the PUF proteins from delaying meiotic entry.

### Repression of PUF proteins by lincRNAs may be evolutionary conserved

The role of RNA in forming RNP phase separated granules is well documented ^58^. Several reports have shown that in mammalian cell cultures the lncRNA *NORAD* binds and restricts two PUF proteins, PUM1 and PUM2 ^35, 36^. Upon DNA damage, the expression of *NORAD* increases. It binds the PUM proteins and shuttles them to phase separated condensates where their ability to repress mRNA is inhibited ^59^. Loss of *NORAD* leads to a significant decrease in the abundance of transcripts that are bound by the PUM proteins, as well as changes in cell cycle and chromosome instability ^59, 60^. These reports on *NORAD* bear similarities to our report on the worm’s lincRNAs. We show here that as cells move out of *C. elegans* gonad niche, the total expression of the lincRNAs that bind a PUF protein increases, and they inhibit FBF-2 action by sequestering it to condensates. Accordingly, deletion of the lincRNA genes affects the cell-cycle and leads to a decrease in the abundance of the transcripts that are bound by the PUF protein. These similarities point towards an evolutionary conserved role for lncRNAs; in mammalian cells, lncRNAs respond to DNA damage and in worms they facilitate correct entry into meiosis. Moreover, PUF proteins are expressed in germline stem cells, not only in *C. elegans*, but also in *D. melanogaster, D. rerio,* mice and humans. Mutations in these genes lead to fertility defects in oogenesis and spermatogenesis ^3, 5, 61–66^. Specifically, these defects include reduction in follicle number in the mature female and establishment of the primordial follicle pool. Interestingly, Pum2 is localized in granules within mice oocytes ^62^. Taken together, many aspects of the PUF-lincRNA system described here are also present in other metazoans. Experiments in mammalian gonads may therefore test if *NORAD,* or other lncRNAs, act to repress PUF proteins during oogenesis in these organisms.

Although the three lincRNAs discussed here are highly expressed in the gonad (*linc-7* abundance is among the top of the transcripts present in the PZ, higher even than most oogenesis genes) ^37^, in comparison with the total number of FBF bound transcripts, they are only a minor fraction. Elguindy et al. have previously reported how a minority of lincRNAs transcripts can restrict the action of PUF proteins. They demonstrated that the number of *NORAD* transcripts is much lower than the entire cohort of PUM binding transcripts. However, once the PUMs are sequestered within the condensates, other interactions keep them sequestered there. *NORAD* can then shuttle back to the cytoplasm to bind more PUMs. It is therefore quite possible that the lincRNAs can restrict FBF-2 in the perinuclear condensates in a non-equimolar mechanism.

Taken together, our work uncovered a layer of control on the mitotic/meiotic decision that is mediated via lincRNAs and the P granules. This novel targeting mechanism adds a new level of interaction between proteins and RNA which attenuates correct balance between proliferation and differentiation. Further studies on the composition of this sub-population of P granules will extend our understanding on how the interaction of proteins and non-coding RNA can promote correct differentiation processes.

## Methods

### Strains and alleles

All strains were cultured under standard conditions at 20 °C unless otherwise specified ^67^. The N2 Bristol strain was utilized as the wild-type background. Worms were grown on NGM plates with *Escherichia coli* OP50 ^67^. All experiments were conducted using adult hermaphrodites 20-24 h after the L4 stage. The following mutations and chromosome rearrangements were used: LGII: *fbf-1(ok91), fbf-2(q738), fbf-2(q932)* (endogenously 3XV5 tagged allele ^23^)*, fbf-2(q704), linc-4(huj25),* LGIII: *mntSi27 fbf-2 (pXW6.26;patcGFP::FBF-2), mntSi28 (pXW6.27; patcGFP::FBF-1)* ^16^*, unc119(ed3)III,* LGIV: *pgl-1(ax3122[pgl1:gfp])* ^68^*, linc-29 (huj19),* LGX: *linc-7(huj9)*.

### Generation of strains

CRISPR-Cas-9 engineering was conducted utilizing the protocol previously described ^69^ with the modifications detailed in ^70^. All engineered mutations were verified by Sanger sequencing.

YBT41: *linc-29(huj19),* was generated using crRNAs listed in Table S3 and was outcrossed five times. This strain carries a full deletion of the gene.

YBT42: *linc-7(huj17)* was generated using crRNAs and ssODN listed in Table S3 and was outcrossed six times. This strain has eight base deletions in the first FBE element TGTATGAT, 565 downstream to the TSS.

YBT51: *linc-29(huj19)*; *linc-7(huj9)*, YBT79: *linc-4(huj25)*; *linc-7(huj9);* YBT80: *linc-4(huj25)*; *linc-29(huj19),* and YBT81: *linc-4(huj25)*; *linc-29(huj19); linc-7(huj9)* were created by crossing the outcrossed single and double mutants. YBT112: *linc-4(huj25); linc-29(huj19); linc-7(huj17)* was created by crossing YBT80 with YBT42.

YBT88: *fbf-1(ok91) linc-4(huj25)*; *linc-29(huj19); linc-7(huj9)* and YBT89: *fbf-2(q378) linc-4(huj25)*; *linc-29(huj19); linc-7(huj9)*, were created by outcrossing YBT81 with JK3022 and JK3101 respectively. The *linc-4* deletion was introduced by crRNA and ssODN listed in Table S3. The *dpy-10* mutation was repaired by crRNA and ssODN listed in Table S3. YBT103: *pgld-1 prom::patcGFP::fbf-2::fbf-2 3’UTR* and YBT105 (*pgld-1 prom::patcGFP::fbf-1::fbf-1 3’UTR)* were generated by crossing UMT382 and UMT392 (kindly provided by Ekaterina Voronina) respectively, with the N2 strain. YBT114: *fbf-2(q932) linc-4(huj25)*; *linc-29(huj19); linc-7(huj9)* was created by crossing JK5842 with YBT-81. The *linc-4* deletion was introduced by crRNA and ssODN listed in Table S3. The *dpy-10* mutation was repaired by crRNA and ssODN listed in Table S3.

### Cytological analysis and immunostaining

DAPI staining and immunostaining of dissected gonads was executed as previously described ^71, 72^. Worms were permeabilized on Superfrost+ slides for 2 min with methanol at −20 °C and fixed for 30 min in a 4% paraformaldehyde in phosphate-buffered saline (PBS). For 10 minutes, the slides were stained with 500 ng/ml DAPI, followed by a destaining in PBS containing 0.1% Tween 20 (PBST). Slides were mounted with Vectashield anti-fading medium (Vector Laboratories).

Primary antibodies were used at the following dilutions: rabbit α-pH3 (D5692, 1:1000; Sigma), mouse α-pgl-1 (OIC1D4, 1:30,000; Developmental Studies Hybridoma Bank), rabbit α-V5-tag (D3H8Q, 1:1000; Cell Signaling). The secondary antibodies used were DyLight 488-goat anti mouse, Cy3-goat anti-rabbit, Cy3-goat anti-mouse (all at 1:500 dilution; Jackson ImmunoResearch Laboratories).

### Single-Molecule RNA in situ Hybridization (smFISH)

Quasar 570 tagged *linc-7* probes for *linc-7* were purchased from Stellaris and diluted 1:10. Labeling was performed as described in ^73^ with additional washing step using wash buffer after the hybridization stage and prior to DAPI staining. Vectashield was used as an anti-fading medium (Vector Laboratories).

### Identification of germline stages

The Leptotene/Zygotene (LZ) region was defined as the first to last rows of the gonad with >2 nuclei with crescent DAPI-stained morphology. The PZ was defined as all nuclei distal to the first LZ row. The distal half of the progenitor zone was defined as “early”, and the proximal half as “late”.

### Imaging and microscopy

Images were acquired using the Olympus IX83 fluorescence microscope system. Optical Z-sections were collected at 0.30- or 0.60-µm increments with the Hamamatsu Orca Flash 4.0 v3 and CellSens Dimension imaging software (Olympus). Pictures were deconvolved using AutoQuant X3 (Media Cybernetics).

### FBF expression level

Expression quantifications were performed as described in ^23^. In short, wholemount complete three JK5842 gonads were stained with V5 antibody and three YBT105 gonads were fixed and stained with DAPI. The gonads were imaged and captured using identical conditions. Using ImageJ, a segmented line (75 mM width) was drawn from the tip to the end of the PZ and along the LZ stage in a plane in which nuclei were present. The fluorescence level of the V5 was measured using the “plot profile” function. For every gonad the number of nuclei rows was counted and averaged. The actual length was divided by the average number of rows and fluorescence values closest to each row position were averaged. Finally, the value for each row was averaged between the different gonads. The graph presented in Fig. 1c depicts combined data from both zones.

### RNA level analysis

RNASeq data of the lincRNAs presented in Fig. S2 were extracted from the analyses reported in ^38^. Early progenitor zone, late progenitor zone, and leptotene/zygotene refer to segments 1-3 in ^38^ respectively.

### RNA-seq

Worms were washed with M9 buffer from NGM plates and 100 µl of Trizol (#15596026; ambion by Life Technologies) was added. Seven cycles of freezing in liquid N2 and thawing at 70°c were applied. RNA was isolated with the Direct-Zol MiniPrep kit (Zymo Research). Three biological replicates were performed for each strain. Sequencing libraries were produced by KAPA standard mRNA-Seq kit (KAPABIOSYSTEMS). Libraries were evaluated by Qubit and TapeStation. Sequencing libraries were constructed with barcodes to allow multiplexing samples on one lane and were processed by Illumina Miseq sequencer following the manufacturer’s protocol. >320 million reads were generated.

### Computational analyses

Reads were aligned to the *C. elegans* genome version WBcel235 using STAR ^74^ version 2.7.9a. Alignment used gene annotations from Ensembl release 105. The indexing stage was done with parameter-genomeSAindexNbases 12. Raw counts per gene were calculated using featureCounts from Subread version 2.0.1. Normalization and differential expression were calculated with Deseq2 ^75^ version 1.34.0. Calculations were done for genes with at least 10 raw counts using default parameters, with applying independent filtering and manually filtering out raw reads of *linc-4, linc-7 and linc-29* for 3xlnc. Genes with an adjusted *p* value below 0.05 and an absolute value of log fold change were considered differentially expressed. For clustering visualizations, DESeq2’s regularized-logarithm transformation (rlog) of the count data was used. Principle Component analysis was done with a function based on plotPCA of Deseq2. MA plot was created using a function based on ggmaplot of ggpubr version 0.6.0, with parameters FDR = 0.05 and FC = 0. Additional annotations were added to the MA plot, using a list of FBF-2 bound-genes ^20^. Heatmaps representing differentially expressed genes were created using ComplexHeatmap ^76^, version 2.10.0 with genes sorted by log fold change and clustered by hierarchical clustering. Annotation for germline enriched genes was created ^39^. GeneOverlap, version 1.30.0 was used to test the significance of the overlap between gene groups. The overlap was tested between upregulated or downregulated genes and FBF-2 bound genes as well as germline enriched downregulated or upregulated genes vs germline enriched FBF2 bound genes, using Fisher’s exact test.

### PZ population and mitotic index analyses

Gonads were stained with DAPI and pH3 Ser10 antibodies. Nuclei in the PZ (see above) were counted manually using ImageJ. Mitotic index was calculated for each gonad as the absolute number of pH3 positive nuclei divided by the total number of nuclei in the PZ population.

### Colocalization of GFP::FBF-2 and PGL-1::GFP with *linc-7* analysis

Gonads of YBT103 and JH3269 were labeled with *linc-7* smFISH probes and images were captured at 0.3 µm for full nuclei volume. To avoid optical aberrations, only the middle third of the nuclei Z stack was analyzed, and the number of foci was then doubled to estimate the number of the entire spherical surface. Foci were determined as colocalized only when a *linc-7* focus was completely colocalized with a protein focus.

### FBF-2 cytoplasm vs perinuclear aggregate analysis

JK5842 and YBT114 wholemount gonads were stained with V5 antibodies and full nuclei were imaged and captured at 0.3 µm Z intervals. Expression of the brightest area of FBF-2 perinuclear foci in each nucleus was measured as well as identical area in the nearby cytoplasm. The value of each cytoplasm measurement was divided by value of the foci, and the relative values for each zone were averaged. The Mann-Whitney U test was used to evaluate significance.

### FBF-2 foci quantification and colocalization with *pgl-1*

JK5842 and YBT114 wholemount gonads were stained with V5 and PGL-1 antibodies and full nuclei were imaged and captured at 0.3 µm Z intervals. To avoid optical aberrations, only the middle third of the nuclei Z stack was analyzed, and the number of foci was then doubled to estimate the number of the entire spherical surface. Foci of FBF-2 which were completely within PGL-1 foci were noted as colocalized.

### Self-progeny quantification

Brood sizes were determined by placing individual L4 worms on seeded NGM plates, transferring each worm to a new plate every 24 h, and counting embryos and hatched progeny across a 3-day period.

### RNAi

To create the *linc-29* feeding vector we used a site-directed mutagenesis PCR method ^77^ to clone the entire *linc-29* cDNA into the L4440 vector between the T7 promotors sites. Feeding RNAi experiments were performed at 20 °C as described previously ^78, 79^. Control worms were fed HT115 bacteria carrying the empty pL4440 vector.

## Acknowledgments

We thank the Caenorhabditis Genetics Center for kindly providing strains. We thank Ekaterina Voronina and Tim Schedl for worm strains. We thank Tzur laboratory members, and especially Kayla Nennig-Kniaz for their helpful comments. We thank Judith Kimble for carefully reading and editing the manuscript. This work was supported by the Israel Science Foundation (#979/21) and by the Ministry of Science & Technology, Israel (#100594) to Y.B.T.

---

**Supplementary Data 1: Differential expression in WT vs mutant strains.** Differential expression in a. *fbf-2* and b. 3xlnc. Columns include general gene details (A-B), raw counts for each sample (C-G), normalized counts (I-N), the DESeq2 calculated baseMean (O), the DESeq2 calculated lfcMLE (Q), the final decision of significance for upregulation in the mutant background (U) and downregulation (V), germline enriched genes as defined in ^39^ (W) and genes bound by FBF-2 as defined in ^20^.

**Supplementary Data 2: List of oligos used in this manuscript.**

