## Supplementary figures and images for "LincRNAs enable germ cells differentiation by promoting PUF proteins condensation"

### Fig. S1

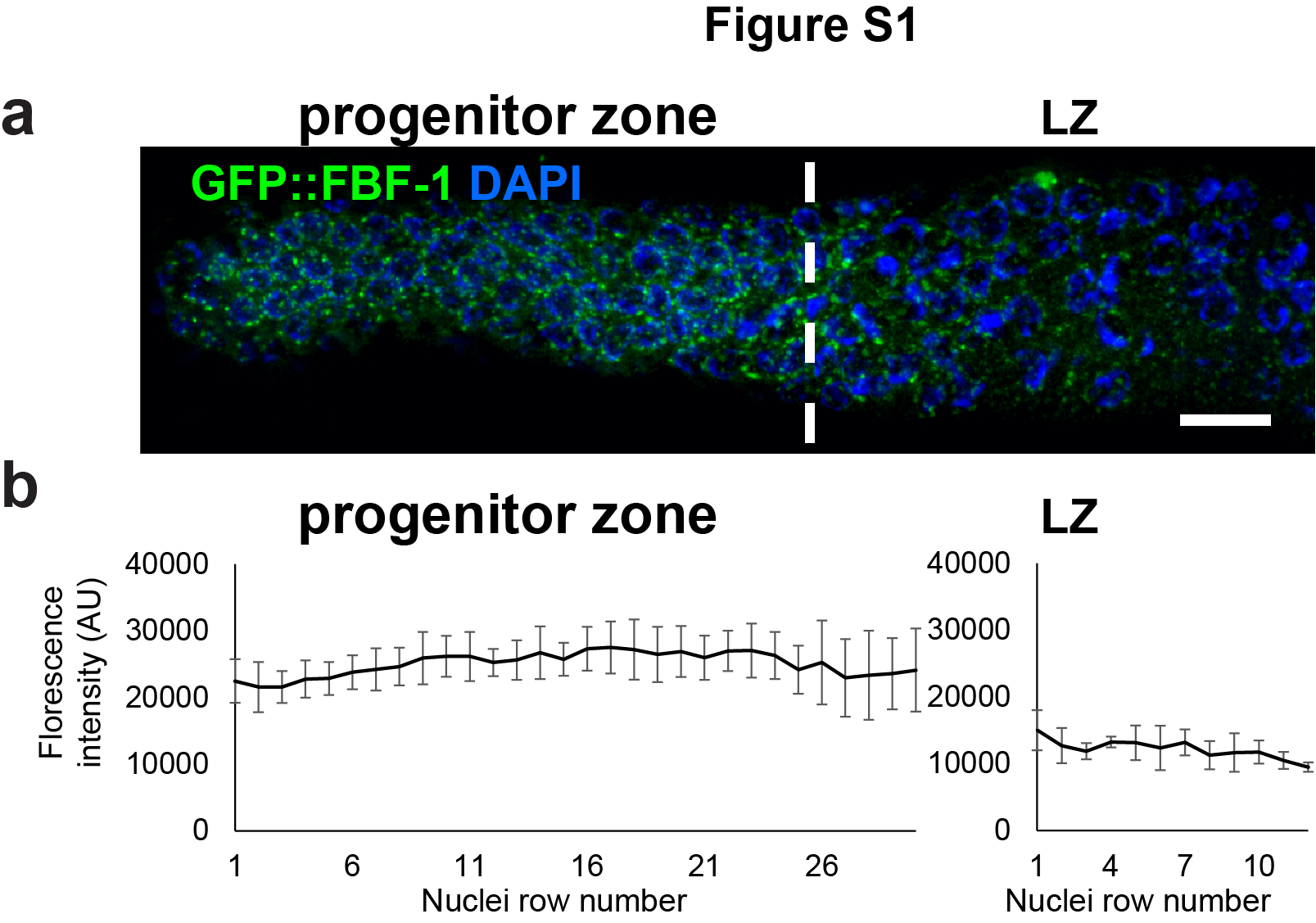

### Fig. S2

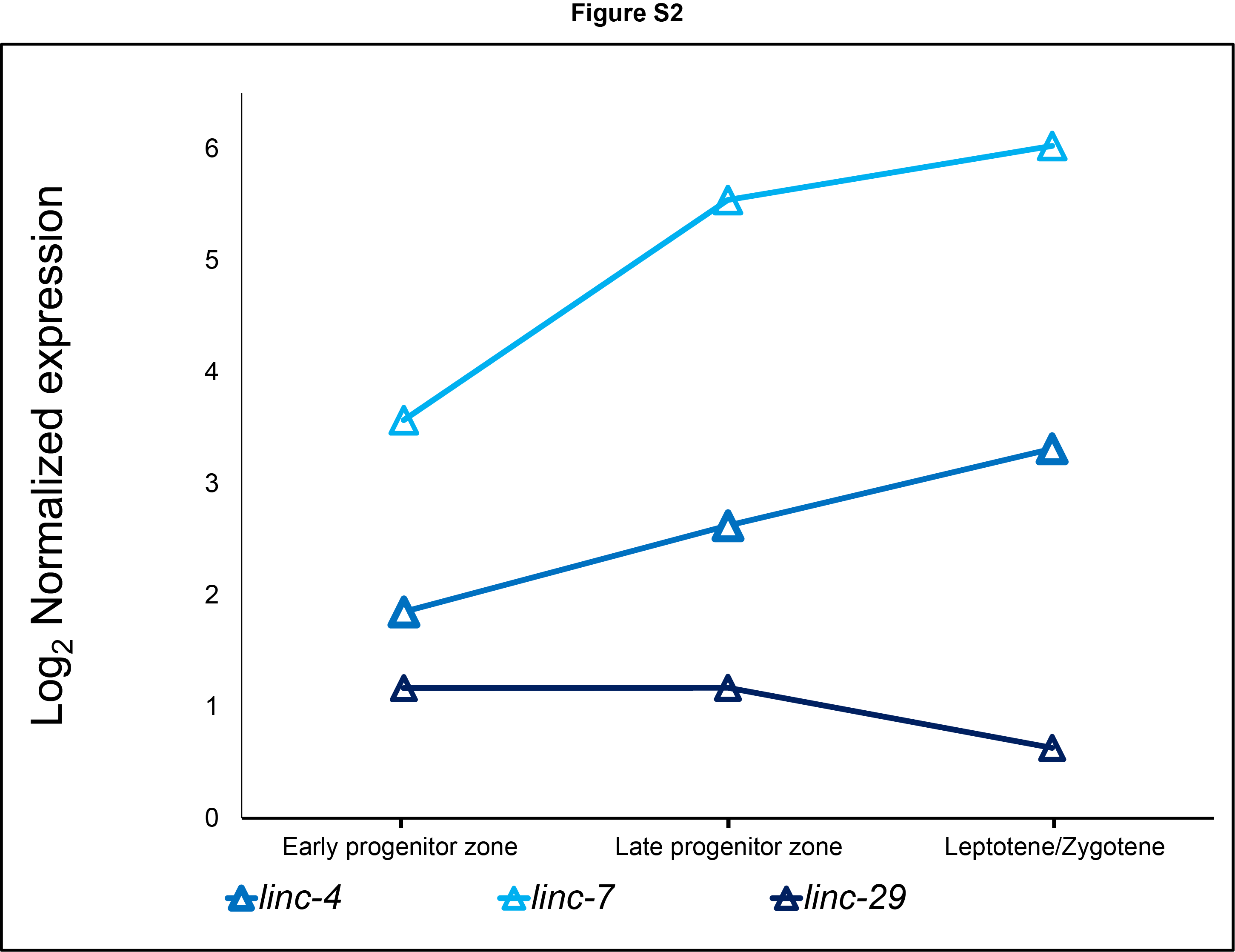

### Fig. S3

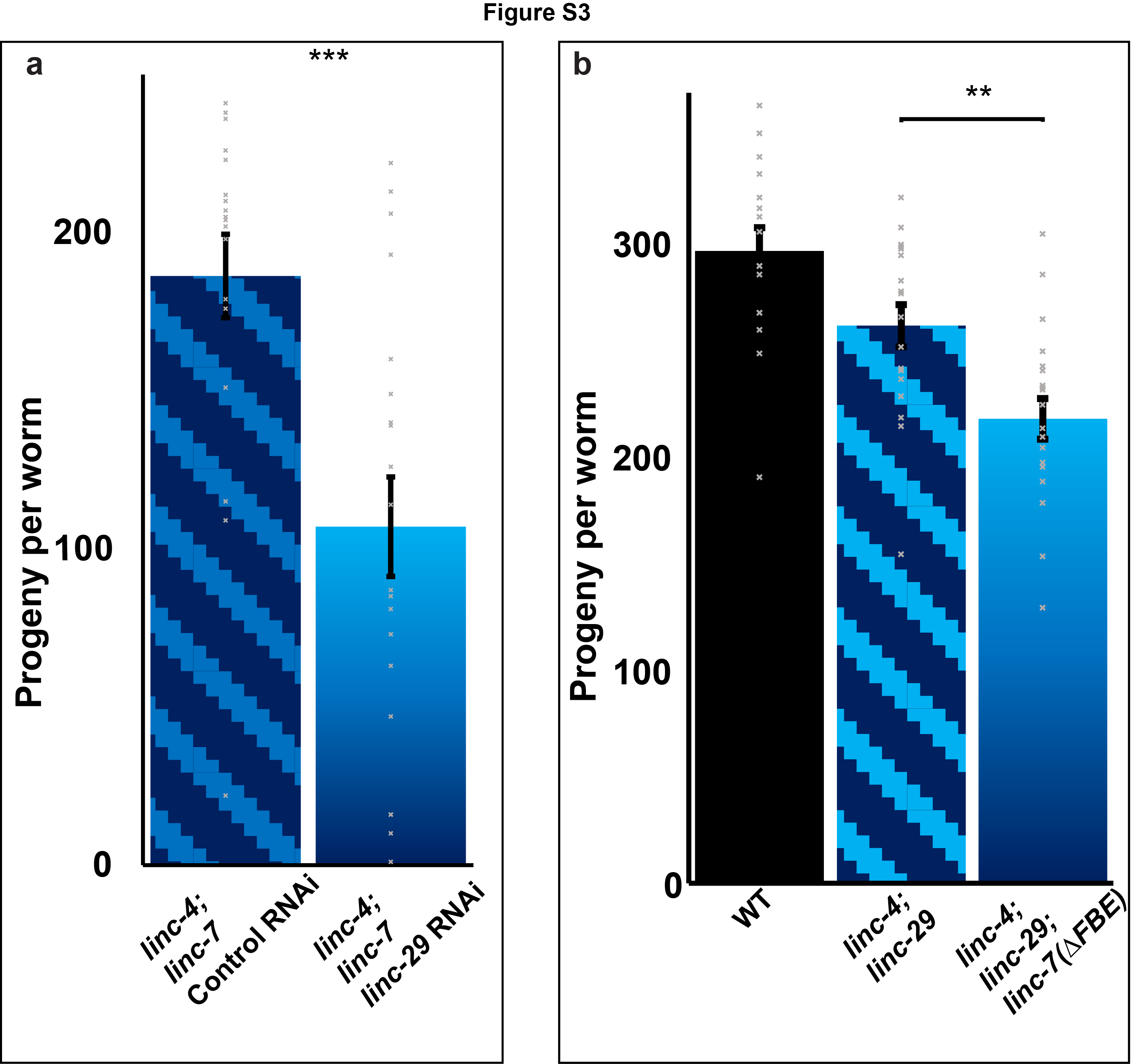

### Fig. S4

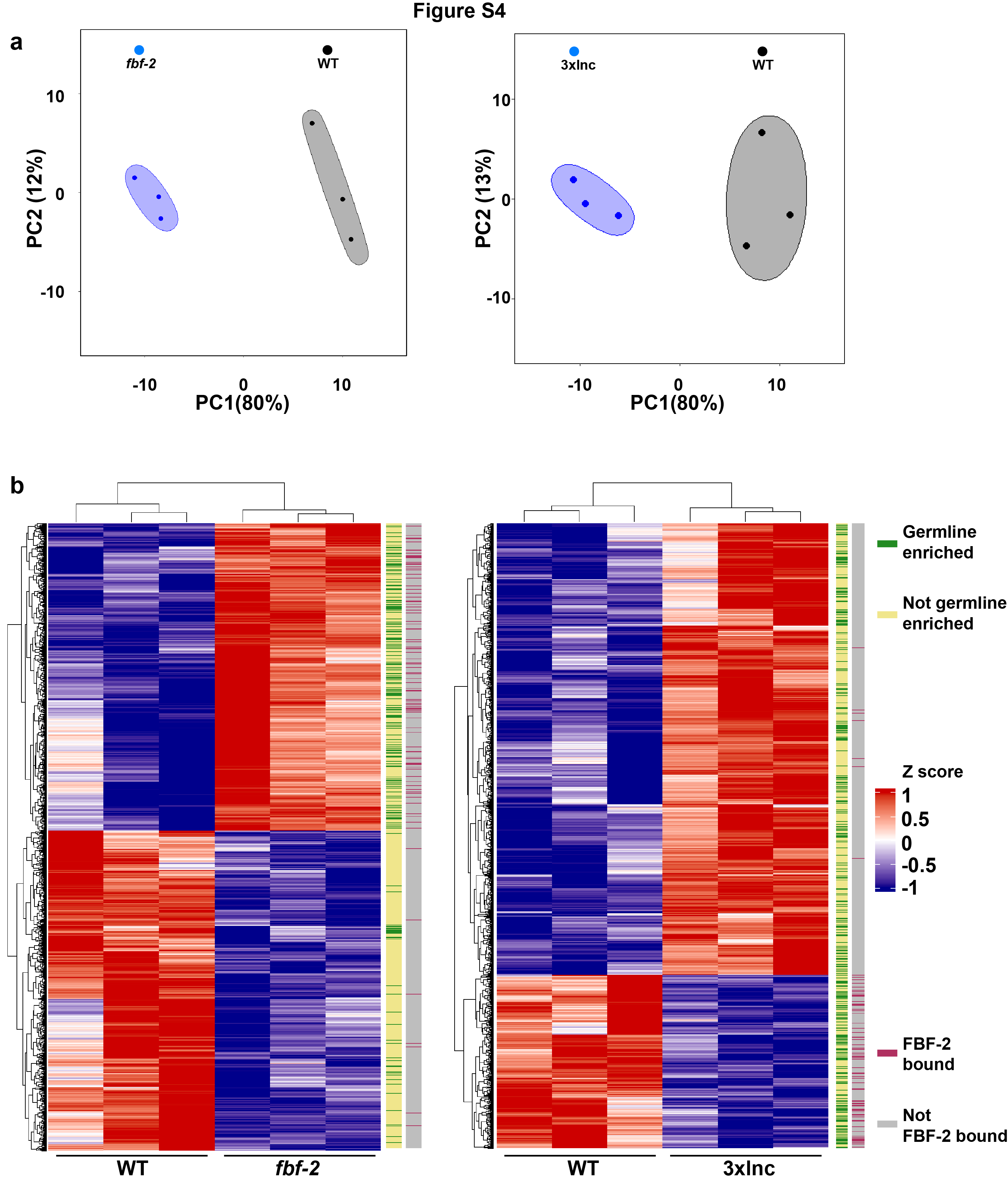

### Fig. S5

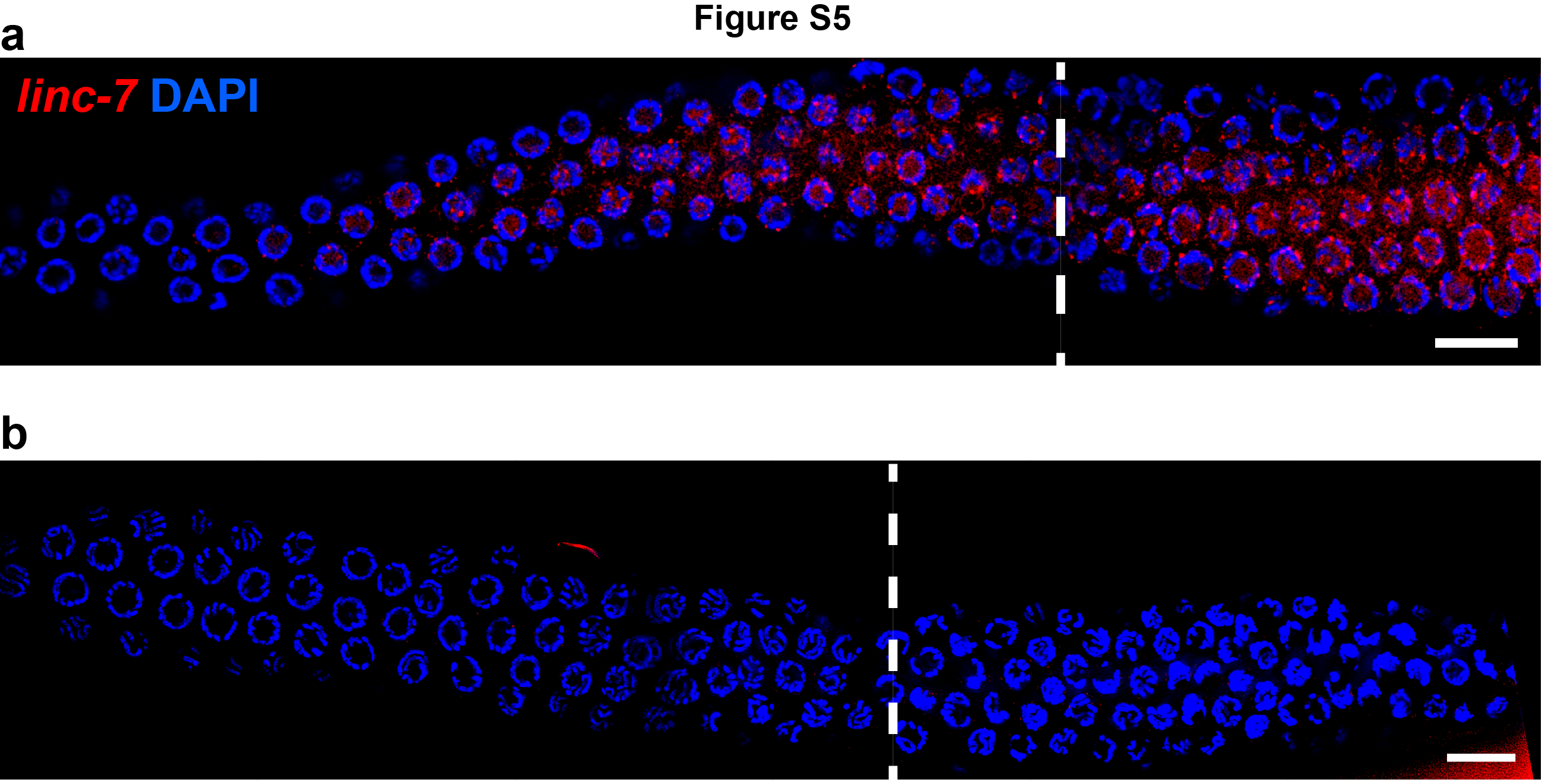
